# UV-B-induced dynamics of constitutive heterochromatin in *Arabidopsis thaliana*

**DOI:** 10.1101/2023.02.17.528949

**Authors:** Philippe Johann to Berens, Kinga Golebiewska, Jackson Peter, Jean Molinier

## Abstract

Sunlight regulates transcriptional programs and triggers the shaping of the genome throughout plant development. Among the different sunlight wavelengths that reach the surface of the Earth, UV-B (280-315 nm) controls the expression of hundreds of genes for the photomorphogenic responses and also induces the formation of photodamage that interfere with genome integrity and transcriptional programs. The combination of cytogenetics and deep-learning-based analyses allowed determining the location of UV-B-induced photoproducts and quantifying the effects of UV-B irradiation on constitutive heterochromatin content in different Arabidopsis natural variants acclimated to various UV-B regimes. We identified that UV-B-induced photolesions are enriched within chromocenters. Furthermore, we uncovered that UV-B irradiation promotes constitutive heterochromatin dynamics that differs among the Arabidopsis ecotypes having divergent heterochromatin contents. Finally, we identified that the proper restoration of the chromocenter shape, upon DNA repair, relies on the UV-B photoreceptor, UV RESISTANCE LOCUS 8 (UVR8). These findings shed the light on the effect of UV-B exposure and perception in heterochromatin dynamics and structure in *Arabidopsis thaliana*.

**Statements and Declarations:** All authors certify that they have no affiliations with or involvement in any organization or entity with any financial interest or non-financial interest in the subject matter or materials discussed in this manuscript.

## Introduction

In plants, the genetic information contained within the nucleus consists of DNA wrapped around a core histone octamer, referred as chromatin, which is organized into discrete chromosomes [1]. Chromosomes can be subdivided in 3 main regions: telomeres, (peri-)centromeres and chromosome-arms with different levels of compaction and containing various genetic elements. Indeed, protein coding genes (PCG) are mainly located in chromosome arms, whilst repeats and transposable elements (TE) are found in telomeric and (peri)centromeric regions [1]. Importantly, chromatin structure organizes the genome into transcriptionally active euchromatin and transcriptionally silenced heterochromatin [2].

During plant development and exposure to environmental cues, chromatin remodeling enables transcriptional activation and/or repression [3–5]. Given that plants use the beneficial effect of sunlight for photosynthesis and for controlling particular developmental programs, many light-dependent mechanisms modulate chromatin shape and thus transcription [5–7]. Notably, factors of the light perception and signaling pathways regulate the level of heterochromatin compaction during different stages of plant development [8]. For example, in Arabidopsis, the photomorphogenesis repressors COP1 (CONSTITUTIVE PHOTOMORPHOGENIC 1) and DET1 (DE-ETIOLATED 1) prevent heterochromatin compaction in etiolated cotyledons [9]. The blue light sensing photoreceptors, CRYPTOCHROMES 1 and 2 (CRY1 and CRY2), are important for the formation of constitutive heterochromatin during the dark-light transition throughout germination [9]. Interestingly, the UV-B photoreceptor UVR8 (UV RESISTANCE LOCUS 8) inhibits the activity of the DNA methyltransferase, DMR2 (DOMAINS REARRANGED METHYLTRANSFERASE 2), leading to the release of silencing of several genomic regions [10] in line with the transcriptional activation and the transposition of the maize TE *Mutator* (Mu) upon UV-B exposure [11, 12]. These studies emphasize that sunlight, the perception of particular wavelengths and the associated signaling pathways play important roles in the regulation of constitutive heterochromatin formation, architecture and silencing, through interconnected mechanisms. Notably, it remains to be documented whether UV-B exposure remodels heterochromatin and to which extent UVR8 could be involved in such dynamic process.

Arabidopsis natural variants, also called ecotypes, originate from different ecological niches characterized by particular environmental features [13]. The different ecotypes offer a wide range of genetic diversity and epigenetic variations allowing to explore the interplay between genome shape and environmental cues. Light intensity, including UV-B regime, strongly vary among the different ecological niches [13]. Several studies, revealed robust correlations between light perception, light intensity and chromocenter shape [14, 15]. Moreover, the heterochromatin content was shown to vary between Arabidopsis ecotypes [16] suggesting the existence of a correlation between chromatin structure and environmental cues. This includes light regimes and likely the damaging effect of particular sunlight wavelength. Indeed, plants have to cope with the deleterious effect of Ultra-Violet (UV). Both UV-A (315-380 nm) and UV-B (280-315 nm) reach the surface of the Earth and lead to the formation of DNA damage affecting genome integrity. While UV-A predominantly gives rise to the formation of oxidatively-induced DNA lesions (8-oxo-7,8-dehydroguanine: 8-oxoG; [17], UV-B is absorbed by DNA bases and directly produces bulky DNA lesions also called photolesions [18]. The 2 main types of photolesions are Cyclobutane Pyrimidine Dimers (CPDs) and 6,4 Photoproducts (6,4 PP). These UV-B-induced DNA lesions are formed between pyrimidines (TT, CC, TC, and CT) leading to DNA helix distortion and interfering with DNA replication and transcription [19].

In plants, photodamage are preferentially repaired by a light-dependent error-free mechanism involving photolyases [20]. In addition, a light-independent process, called Nucleotide Excision Repair (NER) removes UV-induced DNA lesions via 2 sub-pathways: the Transcription-Coupled Repair (TCR) and the Global Genome Repair (GGR) processing photolesions in transcriptionally active and inactive genomic regions, respectively [20]. The existence of these 2 pathways highlights that the epigenomic landscape governs the choice of the repair mechanisms to remove photodamage. Indeed, the NER pathway follows the Access-Repair-Restore model [21, 22] that considers the compaction level of chromatin for the repair kinetics and the mechanisms activated within euchromatic regions (relaxed chromatin) *vs* heterochromatin regions (compacted chromatin; [23]). In addition to the mobilization of particular photodamage repair processes, the genomic regions where photolesions are formed are suspected to be influenced by their epigenomic landscape [24, 25]. Indeed, Rochette et al. [26] reported that di-pyrimidines containing a methylated cytosine (CT, TC and CC) are more prone to form photolesions, suggesting that heterochromatin would likely be more reactive to be photodamaged. However, little is known about genome UV-damageability, albeit the genome*-*wide map of CPD in human cells revealed their preferential enrichment at active transcription factor binding sites [27]. Therefore, more and more lines of evidence support the idea that genome structure, DNA damageability and the photodamage repair choice are interconnected and that environmental cues (*i*.*e*. UV-B regime) may have contributed to shape genomes [28].

In this study, the use of cytogenetics combined with deep-learning-based analyses, allowed documenting the location of UV-B-induced photoproducts and the effects of UV-B irradiation on constitutive heterochromatin shape and dynamics in different Arabidopsis accessions. We found that heterochromatin content, in interphase nuclei of different Arabidopsis ecotypes, negatively correlates with the UV-B regime of their ecological niches. In addition, we identified that constitutive heterochromatin is enriched in photodamage and that UV-B exposure triggers chromocenters dynamics. The way constitutive heterochromatin reshapes depends on the level of heterochromatin content. This holds true in Col-0 and Cvi Arabidopsis natural variants as well as in inter-ecotype hybrids (Col-0 x Cvi). Interestingly, we also report that the restoration of the chromocenter shape, occurring upon photodamage repair, depends on the UV-B photoreceptor UVR8.

Altogether, our observations pave the way for deciphering the range of molecular mechanisms of UV-B-induced heterochromatin dynamics and shaping in *Arabidopsis thaliana* accessions acclimated to different latitudes and thus UV-B regimes.

## Results

### *Arabidopsis thaliana* ecotypes originating from various UV-B regimes exhibit different constitutive heterochromatin contents

In order to determine a putative correlation between UV-B regime and chromocenter shape, we choose four different *A. thaliana* natural variants originating from representative ranges of natural UV-B regimes [29] (Fig. 1a). Ms-0 originates from Moscow, (Latitude 55.75°) and is used as representative of low UV-B exposure, with a mean annual dose of 1418 J/m^2^/day. For high UV-B regime, we used Can-0 from the Canary Islands (Latitude 29.21°) with 4074 J/m^2^/day and Cvi from Cap-Verde Islands (Latitude 15.11°) with a mean annual dose of 5582 J/m^2^/day (Fig. 1a and 1b). Col-0, from Columbia (Latitude 38.3°), the most common ecotype used in research laboratories, serves as control with a mean dose of 2721 J/m^2^/day (Fig. 1b).

**Figure 1.**
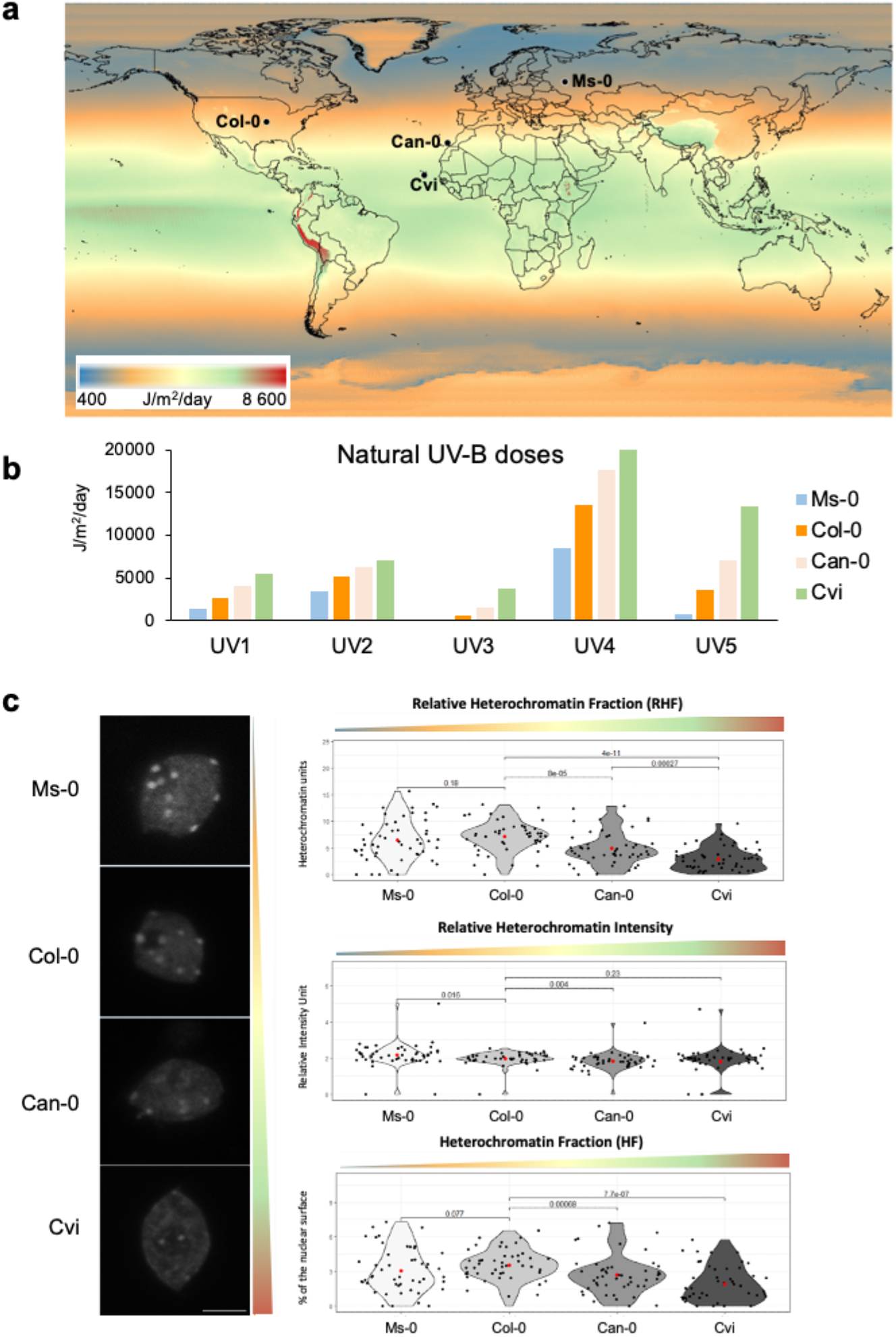
Heterochromatin content of Arabidopsis natural variants originating from different UV-B regimes. **(a)** Worldwide natural UV-B exposure map showing the location of 4 different *Arabidopsis thaliana* ecotypes: Ms-0 (Moscow), Col-0 (Columbia-0), Can-0 (Canary Islands), Cvi (Cape Verde Islands) (adapted from glUV: A global UV-B radiation dataset for macroecological studies [29] **(b)** Histograms displaying UV-B exposure of Ms-0, Col-0, Can-0 and Cvi in their native ecosystem. UV1 = Annual Mean UV-B (in J/m^2^/day); UV2= Mean UV-B of Highest Month (in J/m^2^/day); UV3= Mean UV-B of Lowest Month (in J/m^2^/day); UV4 = Sum of Monthly Mean UV-B during Highest Quarter (in J/m^2^); UV5 = Sum of Monthly Mean UV-B during Lowest Quarter (in J/m^2^) [29]. **(c)** Left panel: microscopy images of DAPI stained Arabidopsis nuclei isolated from Ms-0, Col-0, Can-0, and Cvi leaves. Scale bar = 5μm. Right panel: violin plots showing the distribution of the Relative Heterochromatin Fraction (RHF), Relative Heterochromatin Intensity (RHI) and Heterochromatin Fraction (HF). n= at least 40 nuclei per ecotype. Each black dot represents the measure for 1 nucleus. The red dot shows the mean value. Exact p values are shown (Mann Whitney Wilcoxon test).

According to our working hypothesis, if UV-B regimes have contributed to shape constitutive heterochromatin, we would expect to observe a gradual distribution of the heterochromatin content among the 4 different ecotypes. To test this assumption, we evaluated several chromocenters/nuclei/heterochromatin features in the 4 ecotypes using the Deep-Learning-based tool, Nucl.Eye.D [30] (see experimental procedures for details). As shown in Figure 1c, in interphase nuclei, Relative Heterochromatin Fraction (RHF), Heterochromatin Fraction (HF) and Relative Heterochromatin Intensity (RHI) in Can-0 and Cvi are significantly lower compared to Col-0. Our data are in agreement with the observations of Pavlova et al. [16] reporting that Cvi chromocenters are smaller than those of Col-0. The Ms-0 ecotype, originating from low UV-B regime (Fig. 1a and 1b), exhibits significantly smaller HF compared to Col-0 plants, whilst its RHI is higher than the one measured in Col-0 nuclei (Fig. 1c). Moreover, RHF measurement between Col-0 and Ms-0 does not show a significant difference (Fig. 1c). Thus, it is likely that RHF reaches a maximum from a certain UV-B threshold and/or latitude.

Interestingly, the nucleus size of both Col-0 and Cvi plants do not differ significantly whilst Ms-0 plants exhibit the largest area (Fig. S1a). This observation highlights that the variation of RHF relies mainly on chromocenter size and numbers rather than on the nuclear area (Fig. 1c and S1a). Interestingly, the methylomes of Arabidopsis natural accessions are correlated with geography and climate of origin [31]. Notably, the DNA methylation levels within TEs were positively correlated with latitude [31] as well as chromatin compaction [14]. The Cvi ecotype, that originates from low latitude, displays low DNA methylation level [31]. In addition, it is well established that Arabidopsis plants exhibiting hypomethylated profile (*i*.*e. met1*, defective for *DNA methyltransferase 1* involved in maintenance of CG methylation) have reduced RHF [32]. Thus, the low RHF observed in Cvi plants could partially rely on their low DNA methylation level as well as on the associated structural variations [31]. Hence, modulation of DNA methylation at TE would likely be the consequence of the acclimation to high UV exposure, in agreement with the changes in chromocenter structure induced by variation in light exposure [14].

Although performed with only 4 different ecotypes, these analyses show that RHF negatively correlates with the natural UV-B regime. To further confirm this trend, a similar approach should be enlarged to more Arabidopsis ecotypes originating from various UV-B regimes. Such large scale study could be efficiently set up with the use of Deep-Learning-based tool, Nucl.Eye.D [30]. The identification of ecotypes with different RHF provides useful resources to further characterize the nuclear organization and architecture of Arabidopsis plants acclimated to particular environmental cues (*i*.*e*. light + heat).

### UV-B-induced photodamage are enriched in constitutive heterochromatin

UV-B irradiation predominantly induces CPD and 6,4 PP that are formed between di-pyrimidines [19]. It remains poorly documented whether photodamage are randomly distributed all over the genome or whether they are formed/enriched at particular loci. In order to characterize the location of photolesions, we used the 2 epigenetic divergent ecotypes [31], Col-0 and Cvi displaying high and low RHF, respectively (Fig. 1c). Using fluorescent immunolabeling with anti-CPD antibody, sub-nuclear distribution of CPDs was characterized upon UV-B exposure on 4’,6-Diamidino-2-phenylindol (DAPI) stained Col-0 and Cvi interphase nuclei (Fig. 2a and 2b). As expected, prior UV-B irradiation no immunofluorescent signal was detected, showing the absence or the low level of photodamage formed under our growth conditions (Fig. 2a and 2b). Immediately upon UV-B exposure, CPDs signal became detectable and showed a strong overlap with DAPI labeled chromocenter regions of Col-0 and Cvi ecotypes (Fig. 2a and 2b). A more diffuse signal is present in the nucleoplasm of both ecotypes (Fig. 2a and 2b). Importantly, the immunofluorescent signal intensity is stronger in Col-0 than in Cvi nuclei (Fig. 2a and 2b).

**Figure 2:**
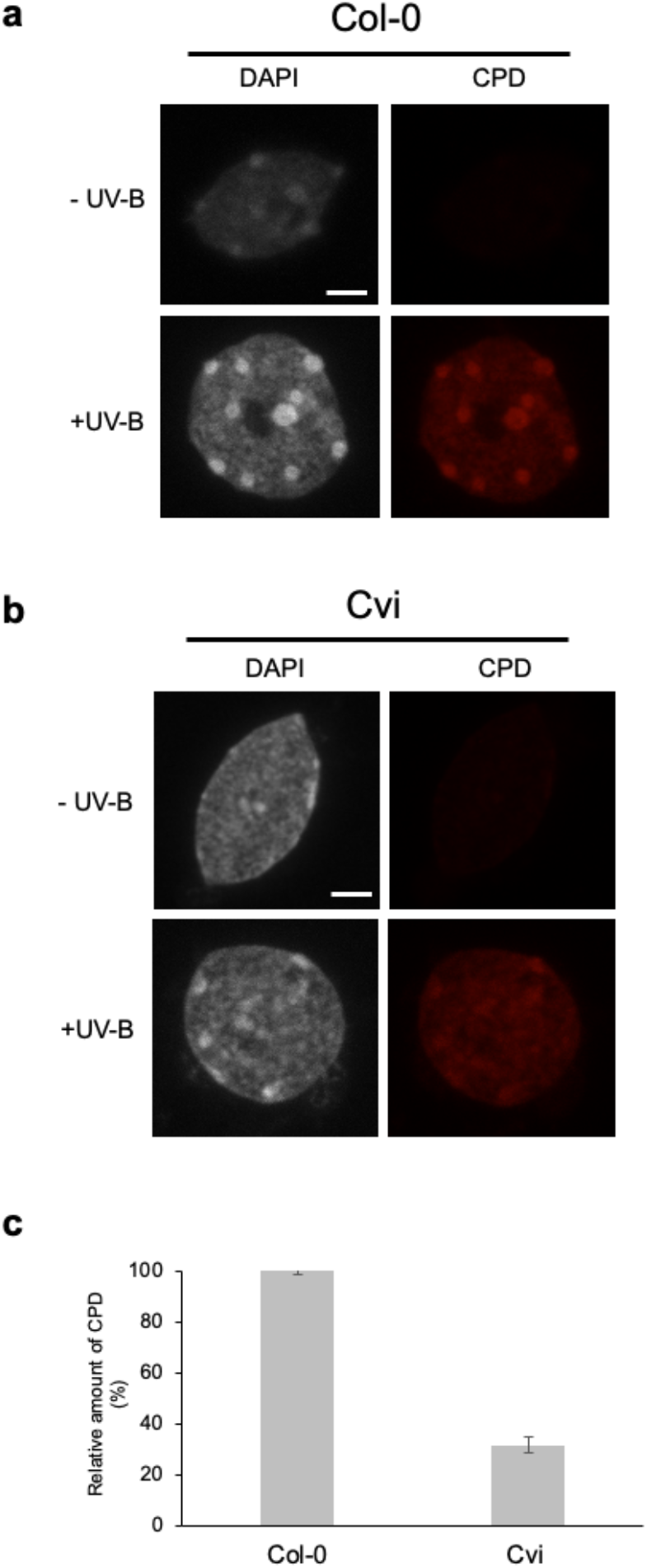
UV-B-induced photodamage localization and quantification. Immunolabeling of CPD on DAPI stained nuclei in **(a)** Col-0 and **(b)** Cvi prepared prior (-UV-B) and immediately upon UV-B exposure (+ UV-B). Scale bar = 5μm. **(c)** Amount of CPD quantified directly after UV-B treatment normalized to the Col-0 plants.

This CPDs’ immunolocalization shows that UV-B-induced photolesions are enriched in constitutive heterochromatin although Col-0 and Cvis’ heterochromatin contents differ significantly. This suggests that, yet unknown, genetic or epigenetic features may facilitate the formation of photodamage in constitutive heterochromatin. Interestingly, di-pyrimidine combinations containing methylated cytosine (CT, TC or CC) are more prone to form a photo-products [26]. In plants, constitutive heterochromatin is enriched in methylated cytosines [33]. Thus, the predominant localization of photolesions in constitutive heterochromatin strengthens the idea that DNA methylation likely contributes to trigger higher reactivity to form photoproducts. In addition to the reduced RHF, Cvi also exhibits low gene body DNA methylation level compared to most of the characterized Arabidopsis natural variants, including Col-0 [31]. Given that DNA photo-damageability could be influenced by the level of DNA methylation, we compared the amount of UV-B-induced CPD in Cvi *vs* Col-0 plants using dot blot. We observed that Cvi plants accumulate 70% less CPDs than Col-0 plants (Fig. 2c). This result is in agreement with the lower immunofluorescent signal observed in Cvi nuclei compared to Col-0 nuclei (Fig. 2a and 2b). Altogether our data suggest that either Cvi developed physiological adaptation to high UV-B irradiance (*i*.*e*. high amount of UV sunscreen) and/or that the hypomethylation profile leads to a low photo-damageability. Indeed, in order to reduce the deleterious effect of UV irradiation plants synthetize UV-absorbing compounds (*i*.*e*. flavonoids), acting as sunscreen protective pigments [34]. Therefore, the combination of particular metabolite profiles together with genetic and chromatin features would likely influence the reactivity of the genome to form photodamage in ecotypes acclimated to different UV-B regime.

### UV-B exposure induces constitutive heterochromatin dynamics

We found that UV-B-induced photodamage are enriched in constitutive heterochromatin of both Col-0 and Cvi ecotypes exhibiting opposite RHF (Fig. 1c). In order to maintain genome integrity, specific DNA repair pathways need to access photolesions for their reversion (DR pathway) or their active removal (NER pathway) [35]. Therefore, constitutive heterochromatin is expected to be remodeled in the first hours following UV-B irradiation to allow photolesions recognition and repair [35]. To analyze the dynamics of chromocenter shape upon UV-B exposure, leaves nuclei of both Col-0 and Cvi *A. thaliana* ecotypes were DAPI stained and analyzed using the Nucl.eye.D script, prior (0) and 2h upon UV-B irradiation. Two hours upon UV-B exposure, the RHF of Col-0 plants significantly decreased whereas RHI remained stable (Fig. 3a and 3b) showing that UV-B irradiation induces heterochromatin dynamics with a transient loss of compaction. Our results are in line with the loss of chromocenter organization and the global rearrangement of the 3D genome observed upon exposure to heat stress, suggesting that various environmental cues lead to the alteration of constitutive heterochromatin shape [4, 36].

**Figure 3:**
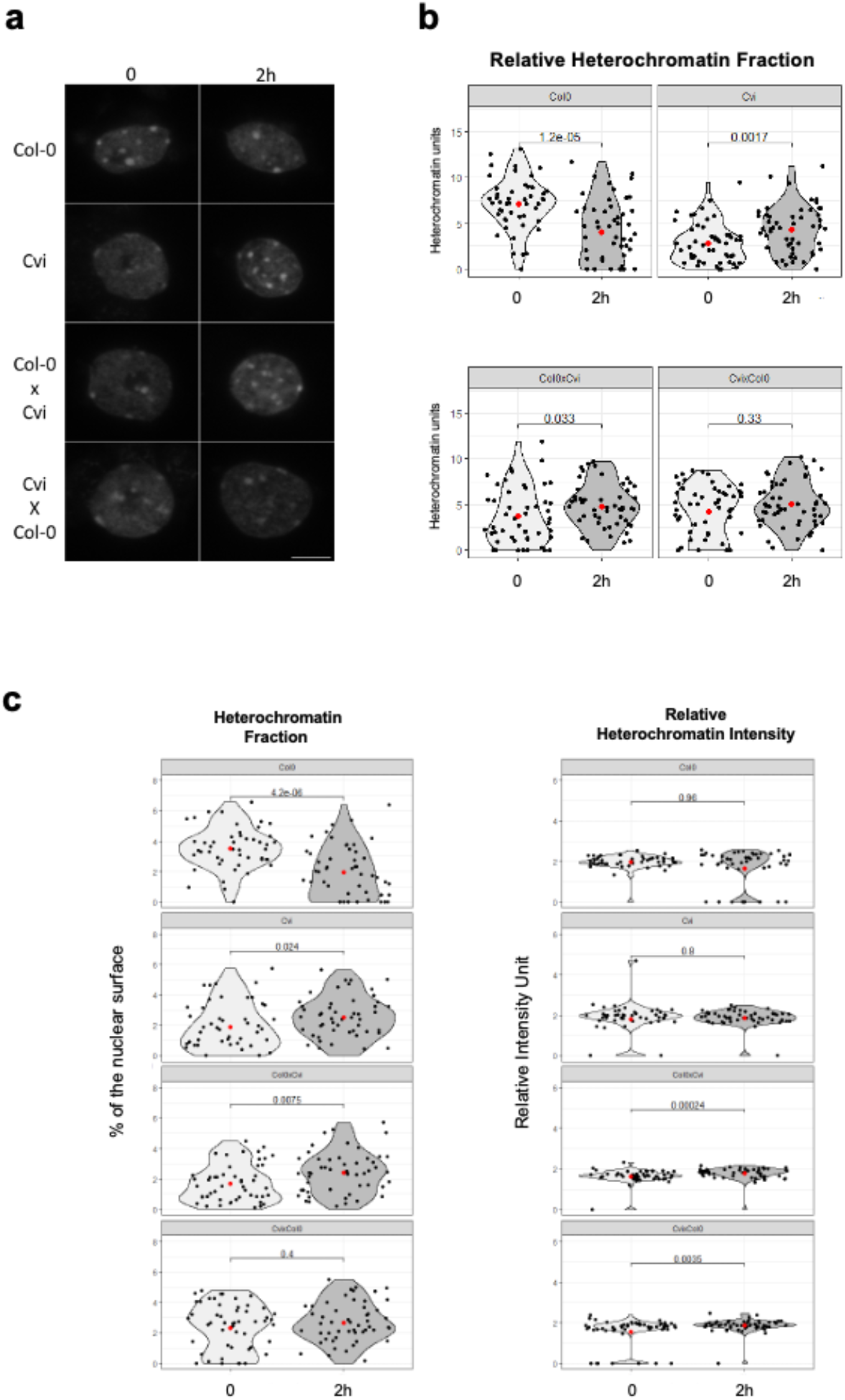
UV-B-induced constitutive heterochromatin dynamics in Col-0, Cvi and Col-0-Cvi hybrid plants. **(a)** Microscopy images of DAPI stained arabidopsis nuclei isolated from Col-0, Cvi, Col-0 x Cvi and Cvi x Col-0 leaves in the control condition (0) or upon UV-B irradiation (2h). Scale bar = 5μm. **(b)** Violin plots illustrating the distribution of the Relative Heterochromatin Fraction (RHF) in a population of at least 45 nuclei per condition as described in **(a)**. Each black dot represents the measure for one nucleus. The red dot shows the mean value. Exact p values are shown (Mann Whitney Wilcoxon test). **(c)** Violin plots showing the distribution of the Heterochromatin Fraction (HF) and of the Relative Heterochromatin Intensity (RHI). Each black dot represents the measure for one nucleus. The red dot shows the mean value. Exact p values are shown (Mann Whitney Wilcoxon test).

In order to test whether UV-B irradiation also induces chromocenter dynamics in Cvi plants, exhibiting low RHF, we used a similar quantitative approach. Surprisingly, Cvi nuclei show a significant RHF increase 2h upon UV-B exposure (Fig. 3a and 3b).

The increase of UV-B-induced RHF measured in Cvi plants is mainly explained by an increased HF and chromocenters number per nucleus (Fig. 3c), arguing in favor of *de novo* heterochromatin formation. Notably, a gain of DNA methylation in heterochromatin was observed in Col-0 plants 24h upon UV-C exposure [37]. Interestingly, the nuclear area of both Col-0 and Cvi plants increases 2h upon UV-B exposure (Fig. S2a) and the number of detected chromocenters changes, with a reduction in Col-0 plants and an enhancement in Cvi plants, in correlation with the variation of RHF (Fig. S2b). On explanation would be that UV irradiation likely induces silencing mechanisms and reshaping of constitutive heterochromatin to prevent further TE reactivation or deleterious chromosomal rearrangements. Indeed, the Cvi ecotype carries a large proportion of TEs in euchromatin domains [16], that correlates with the decondensed shape of the chromocenters. Thus, the Cvi ecotype may have evolved non-canonical regulatory mechanisms of heterochromatin remodeling (*i*.*e*. heterochromatin *de novo* formation) to cope with recurrent high UV-B exposure of its ecological niche. TE and repeats are tightly controlled by the silencing machinery [33]. Defects in DNA methylation as well as exposure to biotic and abiotic stress trigger heterochromatin relaxation, release of silencing and transcriptional reactivation of many TE and repeats [3, 38–40]. For example, in maize, UV-B induces mobilization of the *Mu* TE [11] and in Arabidopsis, UV-C exposure triggers transcriptional reactivation of the *ONSEN* TE, *5S rDNA* cluster and *180 bp* repeats [37]. Hence, complex interplays between epigenomic landscape and genome organization may exist to efficiently control TE and repeats transcription upon exposure to UV irradiation.

Given that Cvi and Col-0 plants exhibit opposite RHF dynamics, we aimed at investigating the effect of UV-B exposure on the transcriptional reactivation of particular centromeric/pericentromeric repeats. Thus, we followed by RT-qPCR, in a time course following UV-B exposure, the transcripts steady state levels of *5S rRNA* and *180 bp* repeats in Col-0 and Cvi plants. Interestingly, prior UV-B exposure, transcripts levels of *5S rRNA* and *180 bp* repeats in Cvi plants are higher than those in Col-0 plants (Fig. 4a and 4b) in agreement with the low RHF quantified in Cvi plants that likely favors transcription in constitutive heterochromatin.

**Figure 4:**
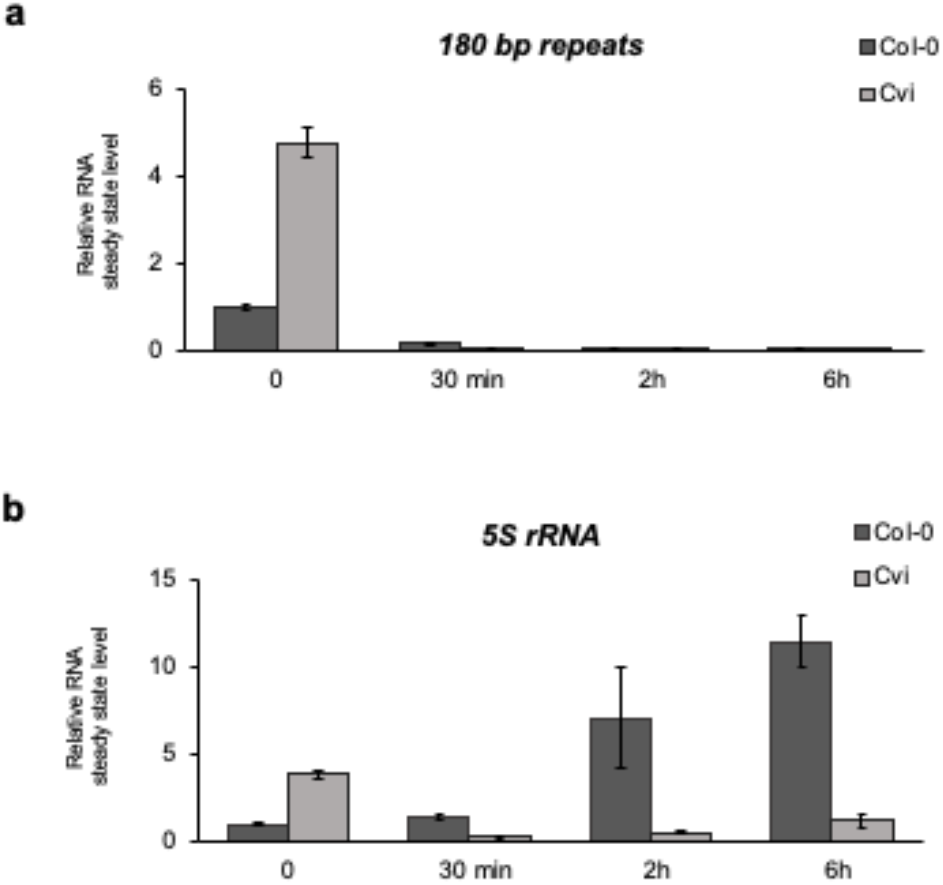
Transcripts levels of *5S* rRNA and 180bp repeats. Transcripts steady state levels of **(a)** *5S rRNA* and **(b)** *180bp* repeats in Col-0 and Cvi plants during a time course following UV-B exposure. RNA steady state levels were normalized to Col-0 (0).

*180 bp* repeats expression profile upon UV-B exposure, shows a reduced transcripts level in both ecotypes (Fig. 4a) suggesting that UV-B wavelength triggers the silencing of these centromeric repeats conversely to UV-C irradiation that releases their silencing [37]. Such different response between the 2 UV wavelengths could be explained by the stronger photodamaging effect of UV-C and by the predominant induction of the DNA Damage Response (DDR), whilst UV-B signaling, in example through UVR8, may act in a parallel pathway.

In Col-0 plants, the *5S rRNA* steady state level gradually increases during the time course (Fig. 4b) correlating with the UV-B-induced heterochromatin relaxation enabling a higher transcriptional activity within pericentromeric regions. The effect of UV-B irradiation on the transcriptional de-repression of the *5S rDNA* cluster, is similar to the one observed upon exposure to heat stress [40] suggesting the existence of common regulatory mechanisms. However, the heat stress-induced release of silencing was shown to be independent of DNA damage signaling pathways [4]. Given that UV-B irradiation leads to the formation of photodamage, predominantly in chromocenters, this scenario would have to be re-evaluated, likely due to the existence of complex interplays between DNA damage, DNA repair, RNA silencing and heterochromatin reshaping [41].

Conversely to Col-0 plants, the amount of *5S rRNA* in Cvi plants exposed to UV-B, decreases throughout the kinetics (Fig. 4b). This correlates with UV-B-induced enhancement of the RHF (Fig. 3b) and thus suggests a reinforcement of the silenced state of this genomic region. Therefore, these observations emphasizes that the low heterochromatin content of the Cvi ecotype, and likely, some particular structural variations, may lead to UV-B-induced heterochromatin *de novo* formation. In addition, the opposite heterochromatin reactivity of Col-0 and Cvi ecotypes highlights that UV-B exposure leads to the mobilization of different molecular processes providing a material of choice to decipher the underlying mechanisms.

### UV-B-induced heterochromatin dynamics is suppressed in Col-Cvi hybrid plants

We found that ecotypes with high and low HRF exhibit divergent heterochromatin dynamics in response to UV-B exposure, highlighting that different mechanisms exist. In order to investigate how different heterochromatin contents and shapes genetically interact together, and react to UV-B exposure, we generated inter-ecotype hybrid plants using parental lines with high (Col-0) and low RHF (Cvi). The crossing was performed in both directions (Col-0 once as mother and once as father), to test a putative parental effect. As shown in Figures 3a and 3b the progenies of both Col-0 ♀ x Cvi ♂ (Hybrid 1: H1) and Cvi ♀ x Col-0 ♂ (Hybrid 2: H2) show an intermediate RHF compared to the Col-0 and Cvi parental lines suggesting that both parents contribute independently and equally to the chromocenter shape in the hybrid plants. In addition, crosses in both directions did not lead to a significant difference in RHF, suggesting that the parental effect is negligeable at this cytogenetic level (Fig. 3a and 3b). Furthermore, the nuclear size as well as the number of chromocenters do not show significant differences (Fig. S2a and 2b).

The inter-ecotype hybrid plants generated between the 2 parents lines exhibiting divergent heterochromatin organization did not lead to major heterochromatin changes. In addition, the plotting of RHI and RHF for each nucleus in H1 and H2 plants, does not highlight the formation of two strikingly different subpopulations, ruling out a sequence specific regulation of the chromocenter structures (Fig. 3b).

To go further in the characterization of chromocenters in these inter-ecotype hybrids, we measured their dynamics 2h following UV-B exposure. In both H1 and H2 hybrids RHF does not vary 2h upon UV-B irradiation (Fig. 3b and 3c) suggesting that independent/antagonist mechanisms, acting in *trans*, likely regulate chromocenters dynamics. Interestingly, the HF significantly increases in H1 plants 2h upon UV-B exposure whereas it remains unchanged in H2 plants (Fig. 3c), highlighting a putative maternal effect originating from the Cvi ecotype. In contrast, the RHI in both H1 and H2 plants displays a similar dynamic, with a significant increase (Fig. 3c). Altogether, these measurements reveal that complex interplays exist to fine tune constitutive heterochromatin in inter-ecotypes hybrids subjected to UV-B irradiation, and that some features are likely under the influence of one parent. Detailed characterization of the epigenetic landscape, chromatin architecture of H1 and H2 plants would allow determining the underlying molecular features contributing to shape heterochromatin in such hybrid plants.

### The UV-B photoreceptor, UVR8, mediates the proper re-establishment of constitutive heterochromatin upon UV-B exposure

We identified that UV-B-induced photodamage are enriched in constitutive heterochromatin and that UV-B exposure triggers chromocenter dynamics. The photoreceptor UVR8 plays a central role in UV-B response, regulating gene expression and photomorphogenesis [42]. In addition, an interconnection has been identified between UVR8 and DNA methylation through the regulation of DRM2 activity [10]. Thus, it was relevant to test whether the UV-B-induced chromocenter dynamics would rely on UV-B perception and thus on UVR8. For this, we exposed WT (Col-0) and *uvr8* plants to UV-B and we measured RHF during a time course. Prior UV-B exposure, RHF of *uvr8* plants does not significantly differ from WT plants (Fig. 5a and 5b). Two hours upon UV-B irradiation, the RHF in *uvr8* nuclei decreases to 8% like in WT plants (Fig. 5a and 5b). Interestingly, 24h upon irradiation, when photodamage are thought to be fully repaired, *uvr8* RHF does not reach the initial level (Fig. 5a and 5b). It remains as low as at 2h whereas in WT plants RHF is back to its initial level measured prior irradiation (Fig. 5a and 5b). These results suggest that the accurate re-establishment of RHF depends on the UVR8 receptor whilst its transient decrease does not. In other words, heterochromatin reconstruction depends on UVR8 while its decompaction does not. To better decipher which nuclear features contribute to the alteration of RHF we also evaluated the HF, the RHI, the nuclear area and the number of chromocenters per nucleus. In WT plants, the drop of RHF at 2h upon irradiation is mainly related to a significant decrease of the HF and chromocenter number per nucleus, whereas the RHI remains stable (Fig. 5c). In *uvr8* plants, the decrease of the RHF, is mainly related to the drop of RHI, 2h and 24 h upon UV-B exposure (Fig. 5c), suggesting that UVR8 regulates predominantly re-establishment of chromocenter structure/organization. Indeed, both Col-0 and *uvr8* nuclear sizes as well as their chromocenter number display the same trends upon UV-B exposure (Fig. S3a and S3b). Therefore, defect in heterochromatin reconstruction observed in *uvr8* plants is mainly explained by impairment of chromocenter reshaping.

**Figure 5:**
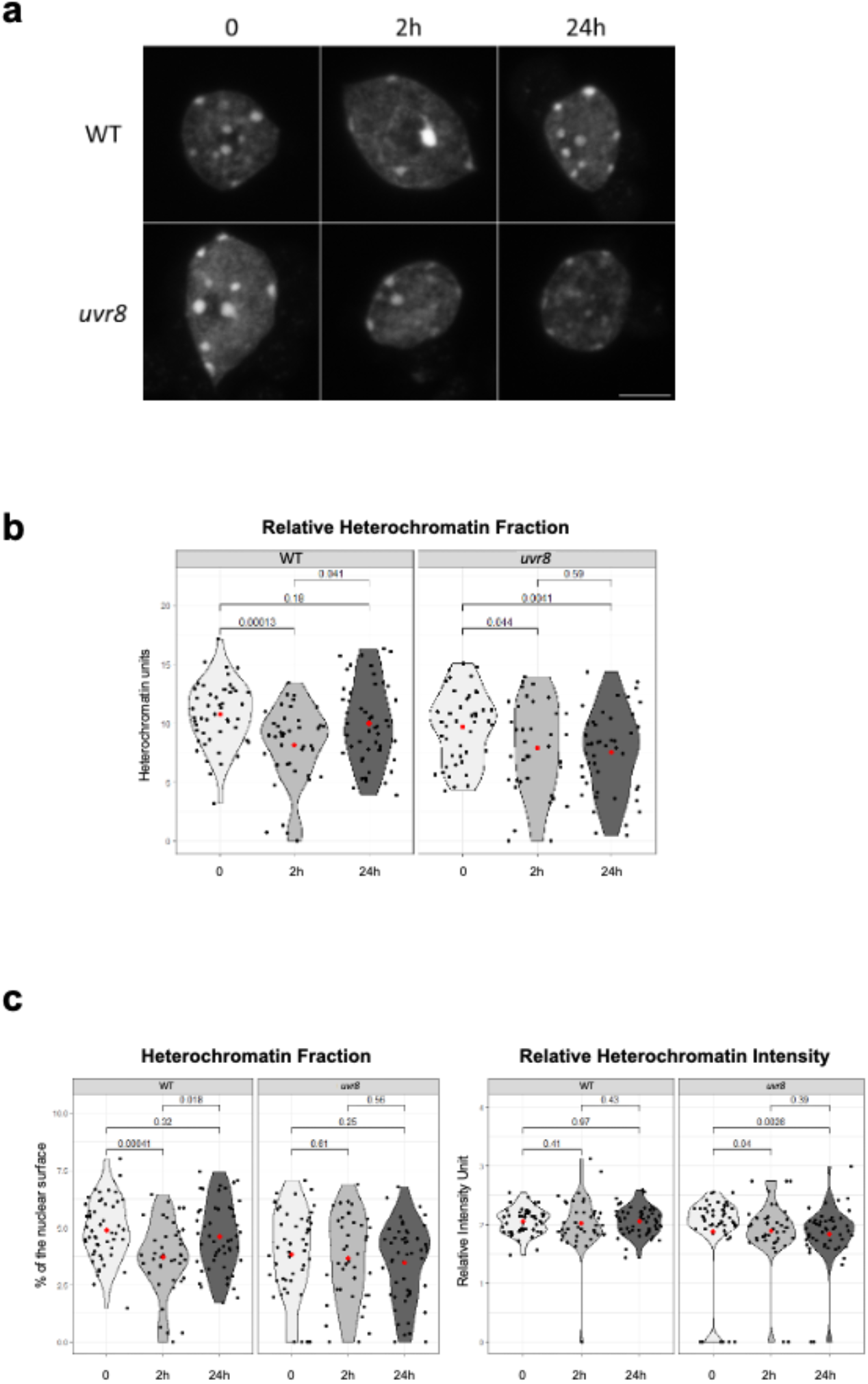
UV-B-induced constitutive heterochromatin dynamics in WT (Col-0) and *uvr8* plants. **(a)** Microscopy images of DAPI stained Arabidopsis nuclei isolated from WT (Col-0) and *uvr8* leaves in the control condition (0) or upon UV-B irradiation (2h, 24h). Scale bar = 5μm. **(b)** Violin plots illustrating the distribution of Relative Heterochromatin Fraction (RHF) in a population of at least 45 nuclei per condition as described in **(a)**. Each black dot represents the measure for one nucleus. The red dot shows the mean value. Exact p values are shown (Mann Whitney Wilcoxon test). **(c)** Violin plots showing the distribution of the Heterochromatin Fraction (HF) and of the Relative Heterochromatin Intensity (RHI). Each black dot represents the measure for one nucleus. The red dot shows the mean value. Exact p values are shown (Mann Whitney Wilcoxon test).

These results demonstrate that UV-B-induced chromocenter dynamics is triggered in an UVR8-independent manner whereas the accurate restoration of chromocenter shape relies on UVR8. Thus, it suggests that photodamage and the associated DNA repair pathways (*e*.*g*. DR and/or GGR) promote heterochromatin relaxation and that UVR8 signaling is likely involved in the regulation of factors acting in heterochromatin reconstruction. Although a recent study revealed a link between UV-B perception and DNA methylation [10] we can rule out that the defect in heterochromatin reconstruction relies on the UVR8-dependent repression of the DNA methyltransferase, DRM2 [10]. Indeed, impairment of DRM2 activity leads to heterochromatin decompaction and thus to smaller chromocenters compared to WT Arabidopsis plants [43]. Hence, we propose that UVR8 would likely preferentially cooperate with DNA damage signaling pathways and/or would mediate activation of, yet unknown, factors involved in re-establishment of the epigenomic landscape and of genome architecture.

## Conclusions

In this study we identified that constitutive heterochromatin content negatively correlates with the latitude where Arabidopsis natural variants originate, suggesting that UV-B regime acts, among other environmental cues, in the shaping of chromocenters. This includes the silencing of TE and repeats which is intimately related to the organization of the epigenetic landscape. Therefore, both genome architecture and epigenome may have co-evolved to specifically shape heterochromatin under the influence of particular environmental factors characterizing the ecological niche of each Arabidopsis natural variant.

Furthermore, we identified that UV-B-induced DNA photodamage are enriched at chromocenters and that a transient remodeling occurs in this area of the chromosome. Importantly, this dynamic differs between Arabidopsis ecotypes exhibiting different heterochromatin contents. Hence, the predominant enrichment of photolesions at chromocenters, as well as their reshaping, underpins the idea that UV-B exposure/regime may have driven their organization/structure together with their remodeling. We also identified a role of the UV-B photoreceptor, UVR8, in the proper re-establishment of chromocenter shape. This highlights that DNA damage signaling, that would preferentially trigger heterochromatin relaxation, is uncoupled from the UV-B signaling process pathway that would rather activate the accurate heterochromatin reconstruction.

Considering our findings, heterochromatin content, shape and dynamics could emerge as a biomarker to reveal UV-B response and plant acclimation to high light exposure.

## Experimental procedures and techniques

### Plant materials and growth conditions

*Arabidopsis thaliana* ecotypes Col-0, Ms-0, Can-0 and Cvi were obtained from the Arabidopsis Biological Resource Stock Center (ABRC, Nottingham, UK). Plants were cultivated *in soil* in a culture chamber under a 16 h light (light intensity ∼150 μmol m^−2^ s^−1^; 21°C) and 8 h dark (19°C) photoperiod. *Arabidopsis thaliana uvr8-6* plants (Col-0 ecotype) were also used [44].

### Ecotypes and UV-B dose regimes

Ecotypes specific longitude and latitude are extracted from https://1001genomes.org/ and used as query for the glUV dataset [29].

### UV-B irradiation

Soil-gown 21-day-old Arabidopsis plants were exposed during 15 min to 4 bulbs of UVB Broadband (Philips - TL 40W/12 RS SLV/25) to deliver a total dose of 6750 J/m^2^. Plant material was harvested prior irradiation for control (0) and during a time course upon irradiation (2h and 24h).

### Tissue fixation, nuclei preparation and immunolocalization of photolesions

Leaves 3 and 4 of soil-grown 21-days old Col-0, Ms-0, Can-0 and Cvi plants are washed 4 times (4°C), at least 5 min, in fixative solution (3:1 ethanol/acetic acid; vol/vol). Leaves nuclei are extracted by chopping fixed tissue in LB-01 Buffer (15 mM Tris-HCl pH 7.5, 2 mM EDTA, 0.5 mM spermine, 80 mM KCl, 29 mM NaCl, 0,1% Triton X-100) with a razor blade. The nuclei containing solution is filtered through 20 µm nylon mesh and centrifugated 1 min (1000 g). Supernatant is spread on poly-lysine slides (Thermo Scientific) and post fixation is performed using a 1:1 acetone / methanol (vol/vol) solution for 2 min. Slides are washed with Phosphate Buffer Saline x1 and incubated for 1h at room temperature in permeabilization buffer (8% BSA, 0.01% Triton-X in Phosphate Buffer Saline x1). For DAPI staining, 15 μl of Fluoromount-G (Southern Biotechnology) with 2 μg/ml DAPI are added as mounting solution before deposing the coverslip.

For immunolocalization of photolesions, leaves 3 and 4 of *in vitro*-grown 21-days old Col-0 and Cvi plants were used. Upon permeabilization slides were incubated over night at 4°C with anti-CPD antibody (Cosmobio) diluted in 1% BSA, Phosphate Buffer Saline x1 buffer. Upon incubation slides were washed at least 3 times with PBS before and secondary antibody coupled to Alexa fluor 488 (diluted in 1% BSA, PBS) was added and incubated for 90 min at room temperature. Finally, slides were again washed 3 times with PBS and 15 μl of Fluoromount-G, with 2 μg/ml DAPI, were added as mounting solution for the coverslip.

### Photodamage quantification

Soil gown 21-day-old Arabidopsis plants (n=40 per ecotype) were irradiated with UV-B (6, 750 J/m^2^). Samples were harvested immediately after irradiation (time 0) and genomic DNA was extracted using plant DNA extraction kit (Macherey-Nagel). CPD content was determined by dot blot as described in [45].

### Microscopy Image acquisition, segmentation and measurements

Image acquisition was entirely performed on a Zeiss LSM 780 confocal microscope using a 64X oil immersion objective. A 405 nm and 488 nm laser excitation wavelengths were used for DAPI, and Alexa Fluor 488/GFP, respectively. Emission DAPI was measured considering wavelengths in the range 410-585. Alexa Fluor 488/GFP emission was measured considering wavelengths in the range 493-630 nm. The same acquisition gain settings were used for all slides of a same experiment. Each image acquisition consists in a Z-stack capture of a 0.64 μm slice distance and the image was reconstructed using the z max plugin of ImageJ.

### Nuclear morphometric parameters measurements using Nucl.Eye.D

The Deep-learning-based tool, Nucl.Eye.D [30] was used to measure the following interphase nuclei morphometric features:

- Number of chromocenters per nucleus
- Nucleus area
- Heterochromatin Fraction (HF): sum of all chromocenters’ areas / nucleus area
- Relative Heterochromatin Intensity (RHI): mean DAPI intensity of chromocenters / mean DAPI intensity of nucleus
- Relative Heterochromatin Fraction (RHF): HF x RHI

### RNA extraction and RT-qPCR

Total RNAs were extracted from 21-day-old soil gown Arabidopsis plants Tri-Reagent (Sigma). Reverse transcription (RT) was performed on 5 μg of total RNA using the cDNA reverse transcription kit (Applied Biosystems) following the manufacturer’s instructions. After RNaseH treatment, 100 ng of purified cDNA were used for quantitative PCR (qPCR). qPCR was performed, including technical triplicates, using a Light Cycler 480 and Light Cycler 480 SYBR green I Master mix (Roche) following manufacturer’s instructions. All primers are listed in Table 1.

**Table 1:**
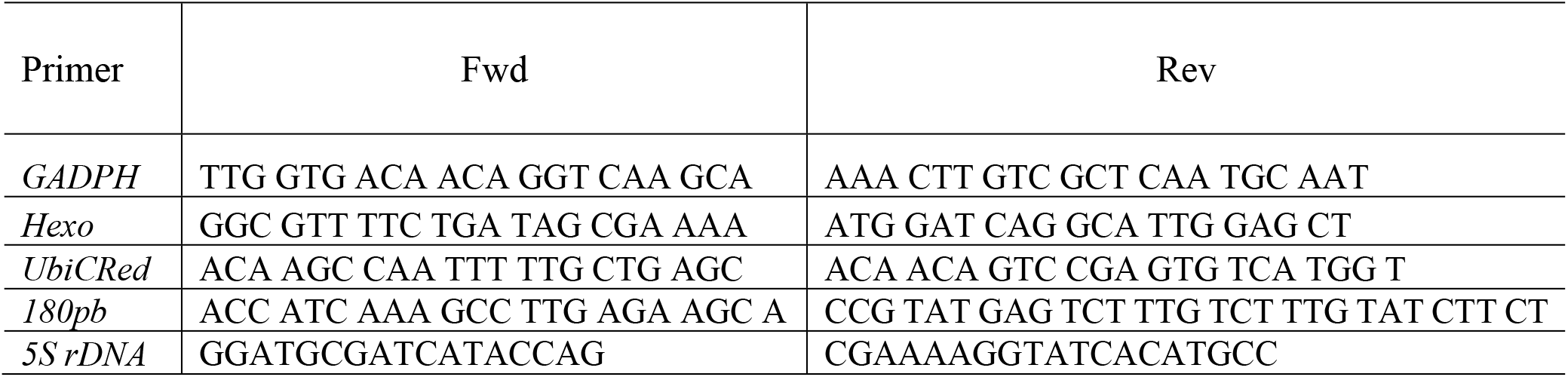
list of primers

### Statistics

Mann-Whitney U or Wilcoxon Matched-Pairs Signed-Ranks tests were used as non-parametric statistical hypothesis tests (http://astatsa.com/WilcoxonTest/). Chi 2 test was used to determine significant difference between categories distribution (**https://goodcalculators.com/chi-square-calculator/**).

## Acknowledgments

We are grateful to Prof. Roman Ulm for providing the *uvr8-6* seeds. This research was funded by a grant from the French National Research Agency (ANR-20-CE20-002) and supported by the EPIPLANT Groupement de Recherche (CNRS, France). K.G. was supported by the ERASMUS program for higher education.

## Supplemental Figures

**Supplemental Figure 1:**
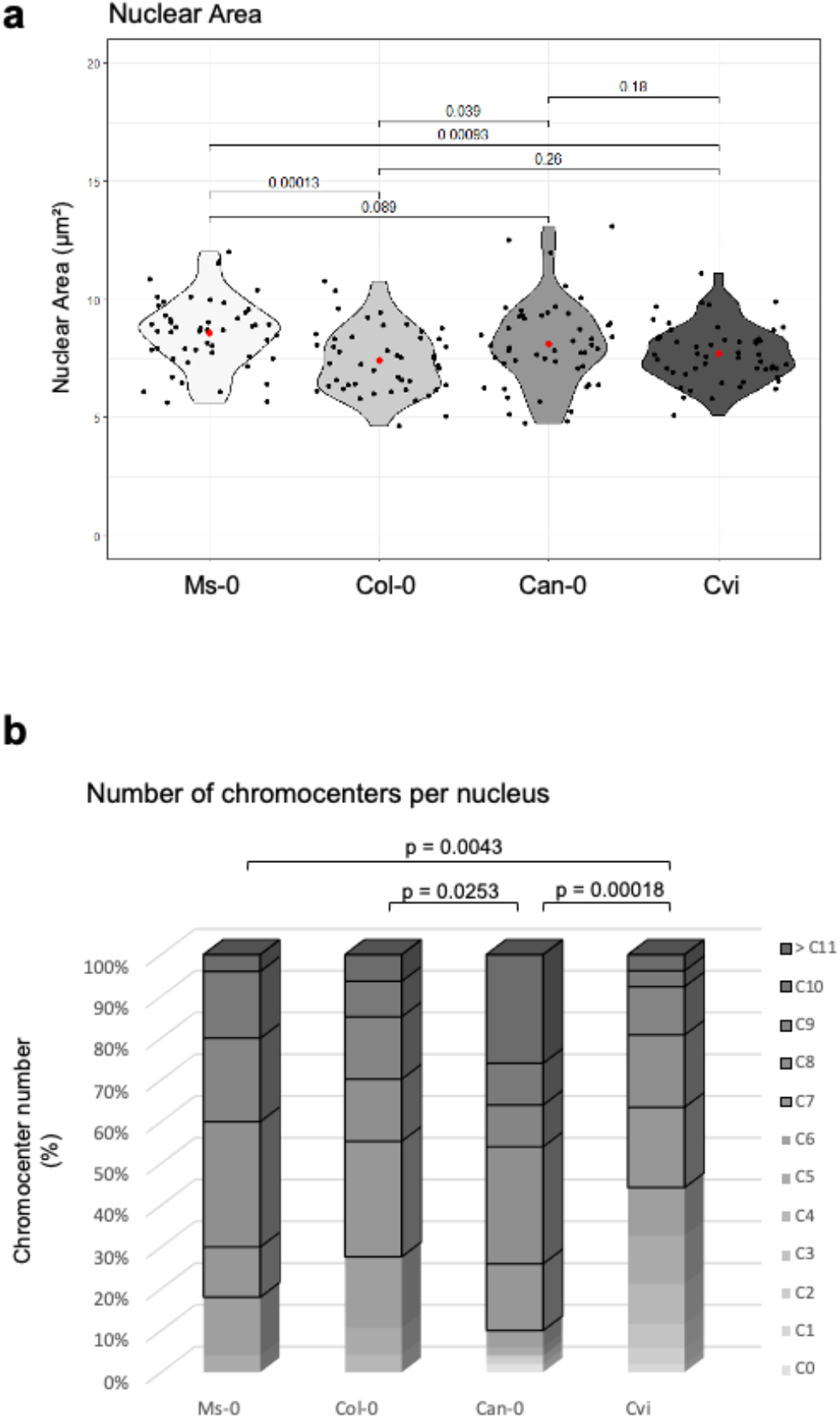
Nuclear area and chromocenter number in Ms-0, Col-0, Can-0 and Cvi ecotypes. **(a)** Violin plots showing the distribution of the nuclear area in Ms-0, Col-0, Can-0 and Cvi plants. Exact p values are shown (Mann Whitney Wilcoxon test). **(b)** Stacked pillar diagram comparing the number of chromocenters per nucleus in in Ms-0, Col-0, Can-0 and Cvi nuclei. Exact p values are shown (Chi-Square test: χ^2^).

**Supplemental Figure 2:**
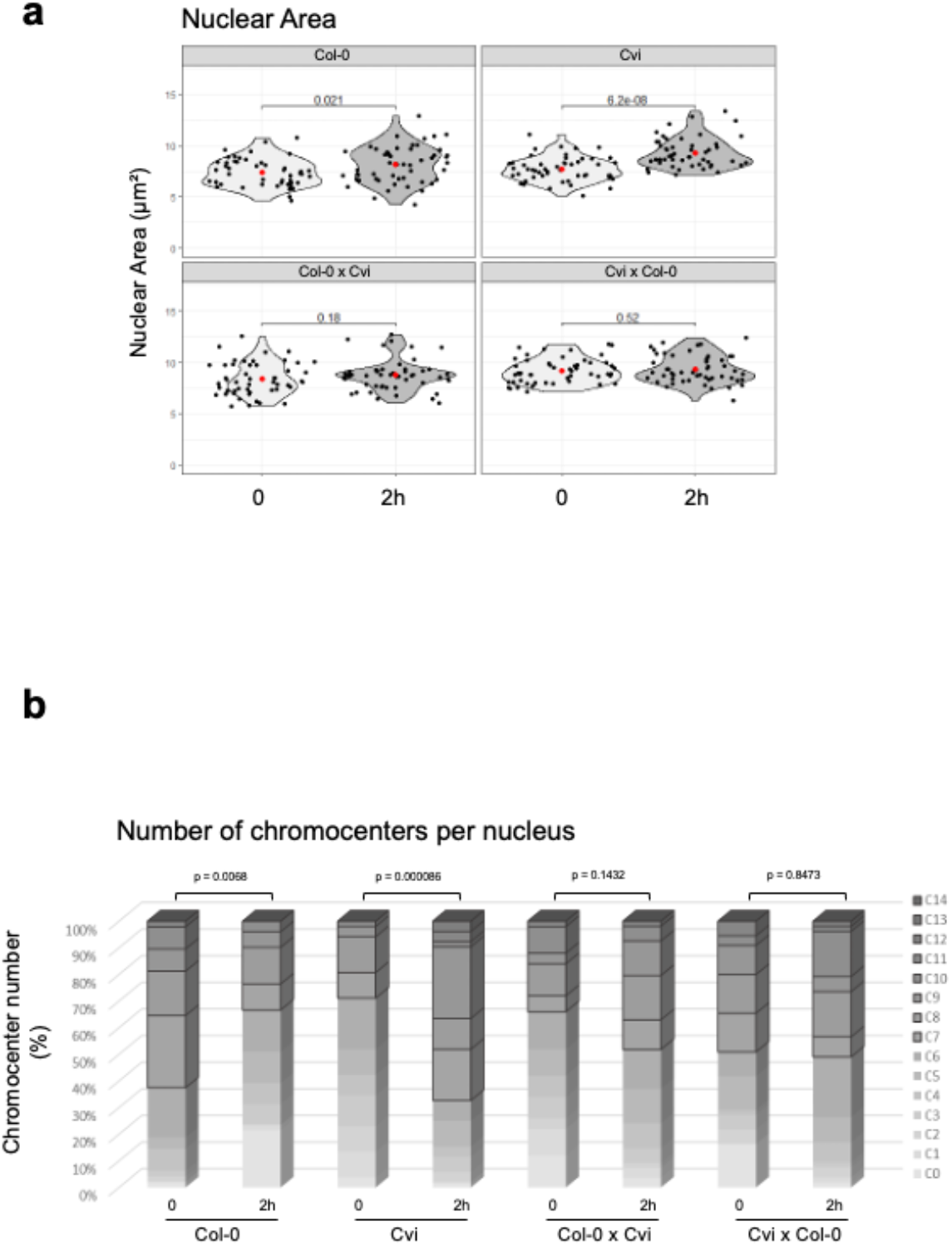
Nuclear area and chromocenter number in Col-0, Cvi, Col-0 x Cvi and in Cvi x Col-0 plants. **(a)** Violin plots showing the distribution of the nuclear area in Col-0, Cvi, Col-0 x Cvi and in Cvi x Col-0 plants before and 2h upon UV-B exposure. Exact p values are shown (Mann Whitney Wilcoxon test). **(b)** Stacked pillar diagram comparing the number of chromocenters per nucleus in Col-0, Cvi, Col-0 x Cvi and in Cvi x Col-0 plants before (0) and 2h upon UV-B exposure. Exact p values are shown (Chi-Square test: χ^2^).

**Supplemental Figure 3:**
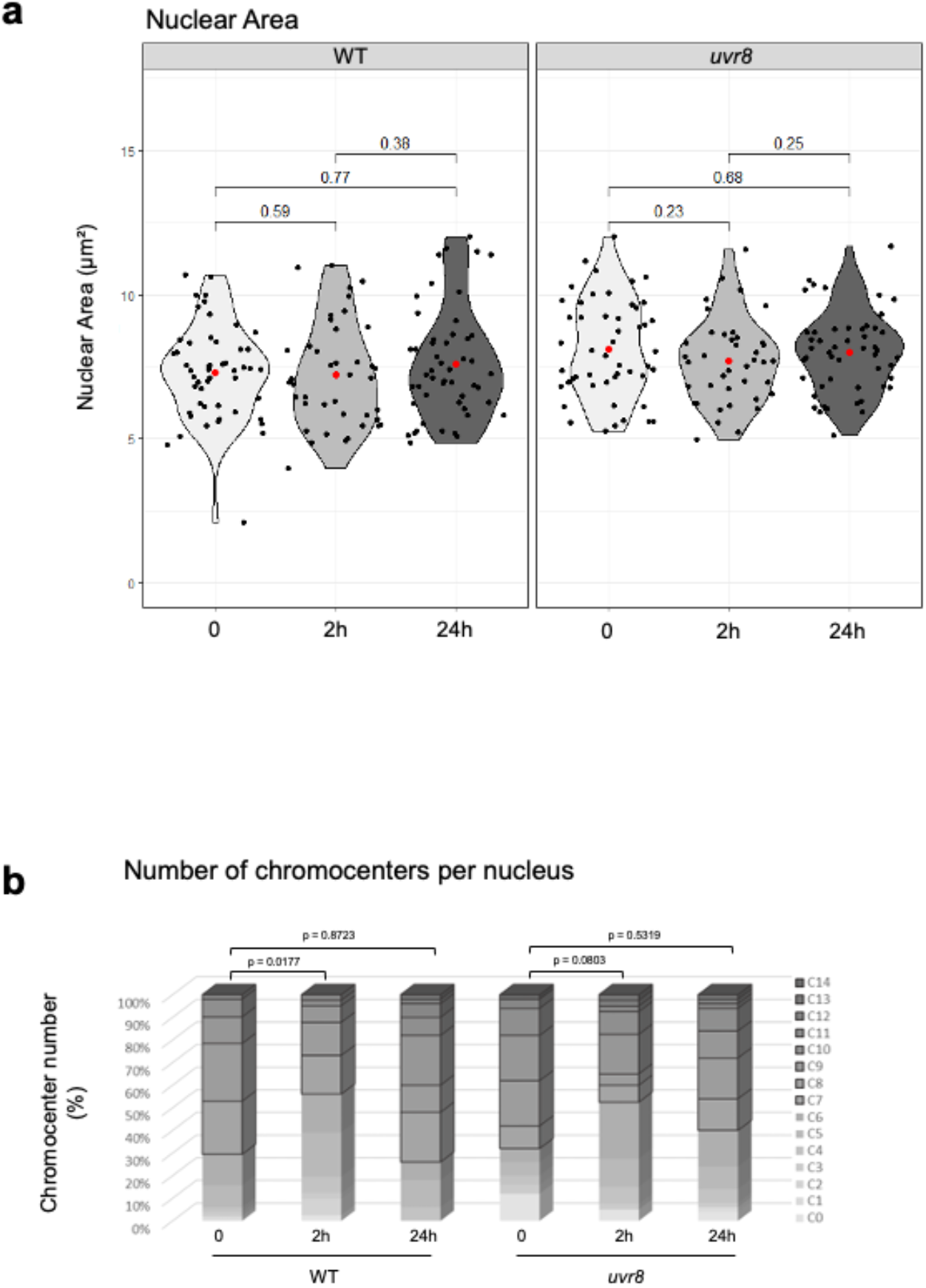
Nuclear area and chromocenter number in WT and *uvr8* plants. **(a)** Violin plots showing the distribution of the nuclear area in WT and *uvr8* plants before (0), 2h and 24h upon UV-B exposure. Exact p values are shown (Mann Whitney Wilcoxon test). **(b)** Stacked pillar diagram comparing the number of chromocenters per nucleus in WT and *uvr8* plants before (0), 2h and 24h upon UV-B exposure. Exact p values are shown (Chi-Square test: χ^2^).

## Notes

### Competing Interest Statement

The authors have declared no competing interest.

